# A Single-Graph Visualization to Reveal Hidden Explainability Patterns of SHAP Feature Interactions in Machine Learning for Biomedical Issues

**DOI:** 10.1101/2025.01.07.631644

**Authors:** Félix Furger, Julien Aligon, Miguel Thomas, Emmanuel Doumard, Cyrille Delpierre, Louis Casteilla, Paul Monsarrat

**Affiliations:** RESTORE Research Center Universit de Toulouse INSERM 1301 CNRS 5070 EFS ENVT France; Universit Toulouse Capitole Institute of Research in Informatics (IRIT) of Toulouse CNRS UMR5505 Toulouse France; CERPOP UMR1295 (EQUITY) Universit P. Sabatier 31000 Toulouse France; Oral Medicine Department and Hospital of Toulouse Toulouse Institute of Oral Medicine and Science CHU de Toulouse France; Artificial and Natural Intelligence Toulouse Institute ANITI France

**Keywords:** Machine learning, Local explanations, Interactions, Patterns in explanations, Biomedical

## Abstract

**Background:** In the last decades, the utility of Machine Learning (ML) in the biomedical domain has been demonstrated repeatedly. Their inherent opacity need augmenting ML with explainability techniques. A common practice in model explainability however, is to focus solely on the explanatory values themselves without accounting for both the main and interaction effects. While this approach simplifies interpretation, it potentially overlooks critical medical information since the nature of the interactions may provide clues to the underlying biological mechanisms.

**Results:** This article introduces a novel method for analyzing explanatory values of machine learning (ML) models, in the form of a comprehensive graphical visualization. The method not only emphasises the individual contributions of the features but also gives insights about the interactions they share with one another. Designed for local additive explanation methods, the proposed tool effectively translates the complex and multidimensional nature of these values into an intuitive single-graph format. It offers a clear window into how feature interactions contribute to the overall prediction of the ML model while aiding in the identification of various interaction types, such as mutual attenuation, positive/negative synergies or dominance of one feature over another.

**Conclusions:** This approach provides insights for generating hypotheses, improving the transparency of ML models, particularly in the context of biology and medicine since living organisms are characterised by a multitude of parameters in complex interactions, a complexity that ensures the “stability” and robustness of structures and functions.

## 1. Introduction

### The need of explainability for the biomedical domain

In the last decades, the utility of Machine Learning (ML) in the biomedical domain has been demonstrated repeatedly [1, 2]. Advanced models, including ensemble approaches and neural networks, have offered high predictive accuracy while leveraging increasingly complex and voluminous datasets. These advancements have significantly contributed to various breakthroughs in diagnostic accuracy and overall healthcare management [3]. However, using such sophisticated models often introduce one major drawback: their inherent opacity, often referred as the “black-box” dilemma [4]. This lack of transparency not only restricts the trust in and adoption of ML but also raises ethical concerns, particularly in sensitive sectors like healthcare, where decision justification for patients and practitioners is crucial [5, 4]. In response to this challenge, emphasis has been placed on augmenting ML with explainability techniques in an attempt to shed light on the reasoning behind the algorithmic decisions [6]. This is all the more important as this explainability can also lead to a better understanding of pathophysiology and the underlying biological mechanisms, even if the inference of causality must be carefully considered.

### Explainability techniques for ML

Explainability techniques can be broadly categorised into local and global methods. Local techniques focus on individual predictions, detailing why a model makes a specific decision, while global techniques provide an overview of the general model behavior. Interestingly, insights gained from local explanations can also be aggregated into global explanations to study both local and global behaviors of the model [6].

Examples of popular local explainability tools for ML are *LIME* (Local Interpretable Model-Agnostic Explanations), *SHAP* (SHapley Additive ex-Planations), and coalition-based methods [6]. Such methods are described as being additive since for a single instance they produce a single vector of weights representing the contribution of each feature (including its interactions with other features) to the prediction. In each case, the contributions approximately sum up to the prediction minus the average prediction for the model. While each method has unique merits [6], *SHAP* stands out as one of the most utilised framework, notably for its ease of use, adaptability to diverse ML methods, and extensive library of functions and visualisation tools.

The intuition behind *SHAP* comes from cooperative game theory and the need for an equitable allocation of “payouts” among “players” using Shapley values. When transposing the idea to the context of ML, the “payouts” and the “players” take the form of the model predictions and the features, respectively. The exhaustive method, referred as the *complete method*, involves evaluating every possible coalition of features, with and without each feature. As coalitions are being evaluated, the impact on the prediction of the presence or absence of a feature in conjunction with other features is used to compute the feature’s contribution [7]. Since the *complete method* is particularly expensive to compute, *SHAP* provides us with a more accessible solution creating perturbations to simulate the absence of a feature. *SHAP* unifies several XAI methods from the literature, in particular *LIME*, a linear local model used to approximate the change in the prediction. The resulting contributions exhibit the previously mentioned additive properties and are referred as SHAP values [7].

For a specific feature and a given sample, the SHAP value represents the additional contribution it provides to the prediction when combined with the set of features used to perform the prediction. By essence, for a given feature and a given prediction, the SHAP value hence encapsulates (i) the main effect, as the individual contribution of the feature when ignoring its interactions with other features and (ii) half the contributions arising from the interactions of the feature with all other features [7].

### Interactions, often forgotten in the explanations

One of the major benefits of ML in the context of biology, and more particularly medicine, is its ability to approximate complex interactions between parameters, regardless of their nature. Indeed, living organisms are characterized by a multitude of parameters interacting in a complex way, a complexity that ensures the “stability” and robustness of structures and functions. A common practice in model explainability however, is to focus solely on the explanatory values themselves (*i*.*e*. SHAP values) without accounting for the decomposition between main and interaction effects. While this approach simplifies interpretation, it potentially overlooks critical medical information since the nature of the interactions (e.g. synergy, mutual attenuation) may provide clues to the underlying biological mechanisms that are not immediately apparent when examining individual predictors.

*SHAP* provides us with ways to extract these interaction effects for some ML models. However, current graphical representations provided by the library are limited. For a model with *n* features, there are *n*(*n* − 1) sets of SHAP interaction values as well as *n* sets of SHAP main effect values to analyze, resulting in a total of *n*^2^ sets of data points to handle. Visualizing each of the *n*^2^ SHAP dependence plots quickly becomes out of reach as *n* increases, hence complicating the task of finding patterns, groupings, and generalising findings for models with many features.

Consequently, this article introduces a novel method for analyzing SHAP interaction values, in the form of a comprehensive graphical visualization. The method not only emphasises the individual contributions of the features but also gives insights about the interactions they share with one another.

## 2. Material and methods

### Illustration context

In order to illustrate the proposed method, the explainable machine learning framework and database to predict physiological aging was considered [8]. This database involves 48 routine laboratory biological variables obtained for 60,322 individuals from the National Health and Nutrition Examination Survey (NHANES) database (1999 - 2018). All details were provided in the dedicated article [8]. As detailed previously, an *XGBoost* model was trained to predict chronological age based on the 48 biological variables as predictors. The SHAP interactions values were subsequently extracted using *TreeExplainer* from the *shap* python library.

### Interaction tensor

In the context of tree-based models (*TreeExplainer, shap* python library), the first layer of interactions can be extracted, corresponding to a 3-dimensional tensor of shape (*n samples, n features, n features*) where *n samples* and *n features* respectively represent the number of instances and features in the dataset. The diagonal entries of the tensor (across the last two dimensions) provide the main effect of each feature on the model prediction for each sample, without accounting for any interactions. The off-diagonal entries represent the interactions between pairs of features. Specifically, the entry at position (*i, j, k*) represents the interaction between feature *j* and feature *k* for sample *i*. These entries capture the combined effect of the two features that can not be attributed to their individual contributions. Such interactions are symmetric, meaning that for sample *i*, the interaction between feature *j* and feature *k* is identical to that between feature *k* and feature *j*.

### Detection of the interaction trends

To inform about whether the interaction between a feature and another tends to amplify or attenuate the main effect of the feature itself, each interaction is multiplied by the sign of the SHAP contribution of the main effect. For instance, when the SHAP interaction value of *Glycohemoglobin* with *Triglycerides* (Fig. 1B) is multiplied by the sign of the SHAP main effect of *Glycohemoglobin* (Fig. 1A), it becomes clear that high values of *Triglycerides* tend to amplify the main (positive) effect of *Glycohemoglobin* (Spearman correlation coefficient *ρ* = 0.78, Fig. 1C, D). In contrast, high values of *Cholesterol* and *ALT* tend to attenuate the main (positive) effects of *Glycohemoglobin* and *Cholesterol*, respectively (*ρ* = − 0.78, *ρ* = − 0.82, Fig. 1A-D).

**Figure 1:**
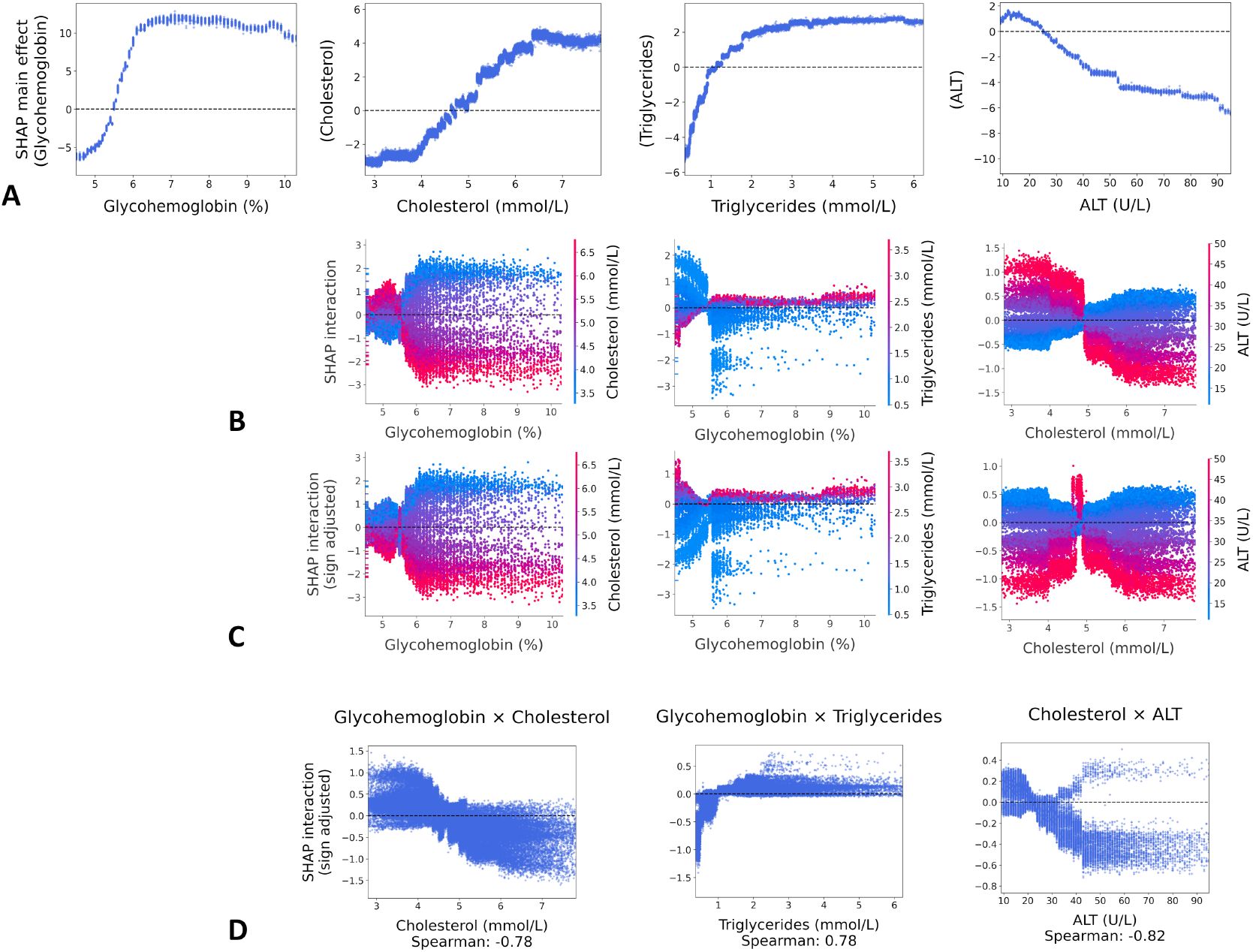
Representation of the treatment process to build the interaction graph. **(A)** Partial dependance plots displaying the SHAP main effect of each variable as a function of its raw value. Here, *Glycohemoglobin, Cholesterol* and *Triglycerides* have positive main effects while *ALT* has a negative main effect. **(B)** Partial dependence plots displaying SHAP interaction values for a pair of features as a function of the first feature’s raw values (x-axis) and the second feature’s raw values (color gradient). **(C)** Same as (C) but displaying SHAP interaction values after their multiplication with the sign of the first feature’s SHAP main effect values from (A). For example, high values of *Cholesterol* tend to attenuate the main effect of *Glycohemoglobin* whereas high values of *Triglycerides* tend to amplify the main effect of *Glycohemoglobin*. **(D)** Detection of the interaction trends through Spearman’s correlation coefficient. *Glycohemoglobin* × *Cholesterol* as well as *Cholesterol* × *ALT* will result in blue arrows on the graph whereas *Glycohemoglobin* × *Triglycerides* will result in a red arrow.

Formally, if *I* is the 3-dimensional interaction tensor returned by *Tree-Explainer*, a new 3-dimensional interaction tensor *S* can be computed so that:

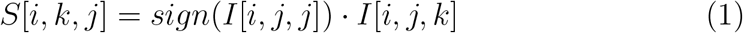

where *I*[*i, j, j*] is the main effect of feature *j* (diagonal entry). The resulting tensor has been transposed along its last 2 dimensions (*S*[*i, k, j*] instead of *S*[*i, j, k*]) so that when feature *j* interacts with feature *k*, it is possible to see whether the values of feature *k* amplify or attenuate the main effect of feature *j*.

Consequently, computing the Spearman correlation coefficient allows to extract the direction and the strength of this association. The Spearman coefficient was chosen for its ability to detect monotonic non-linear relationships and its sign indicates the direction of monotonicity. However, Pearson’s correlation coefficient was also computed for its ability to detect monotonic linear relationships.

## 3. Results

Each node of the graph (Fig. 2) represents a biological variable, i.e. feature. Its size relates to the mean SHAP value of this feature’s main effect while its color relates to the direction of the effect (i.e. red and blue for positive and negative correlation with predicted age, respectively). As an example *Triglycerides, Cholesterol* and *Glycohemoglobin* are red nodes (positive correlation with predicted age) while *ALT* is a blue node (negative correlation with predicted age).

**Figure 2:**
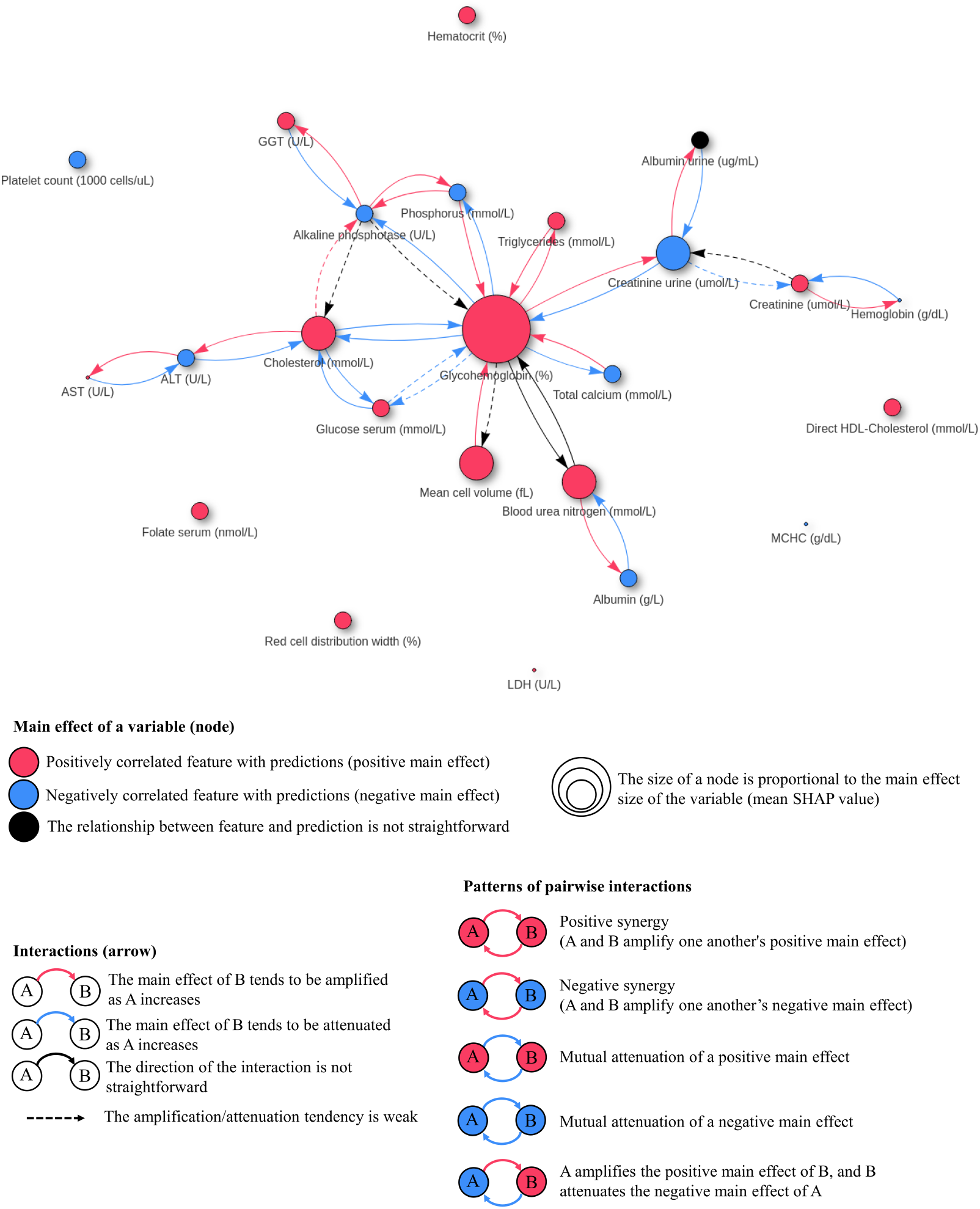
Representation of the interaction graph. Each feature is a node whose size is proportional to the main effect and whose color indicates the sign of the correlation. Arrows between nodes are interaction values whose color indicates amplification or attenuation of the main effect of a feature when the other feature increases. When the correlation is weak (Spearman’s correlation between -0.3 and 0.3), the arrow is rendered dashed. If Spearman’s and Pearson’s coeffic1i1ents are of opposite signs, the corresponding node or arrow is rendered black as a warning. This specific graph was obtained when setting the interaction threshold to 0.22 (after normalising the top 100 interactions, we display only those exceeding 22% of the maximum normalised value).

Each arrow from feature *k* to feature *j* indicates whether an increase in feature *k* amplifies or attenuates the main effect of feature *j*. High values of *Glycohemoglobin* amplify the previously identified positive main effect of *Triglycerides*, resulting in a red arrow on the graph but attenuate the previously identified positive main effect of *Cholesterol*, resulting in a blue arrow on the graph. High values of *Cholesterol* amplify the previously identified negative main effect of *ALT*, resulting in a red arrow on the graph. Additionally, a threshold can be set on the mean absolute SHAP values of nodes and arrows to allow only those of a certain importance to be displayed. In contrast, it is possible to observe situations in which the mean absolute SHAP value is high but the choice of color for the node or arrow is not straightforward. When Spearman’s coefficient is between -0.3 and 0.3 (default value, parameter of the function), the correlation is considered weak, the arrow is rendered dashed and particular attention must be paid to the visual analysis. In addition if Spearman’s and Pearson’s coefficients are of opposite signs, the corresponding node or arrow is rendered black as a warning. Such situations suggest a careful graphical analysis of the relationship is necessary.

## 4. Discussion and conclusion

This proposal shows the benefit of dissociating raw contributions from interaction contributions in additive explanation models in machine learning. This optimized visualization reveals the central role of *Glycohemoglobin*, that not only directly strongly impacts the prediction but is in interaction with a large number of features. The patterns of pairwise interactions demonstrate a dominant effect of *Glycohemoglobin* for several interactions as well a positive synergy with *Triglycerides* and a mutual attenuation with *Cholesterol*. Such mutual attenuation may reveal homeostatic regulatory mechanisms.

Even if this visualization only highlights monotonic trends, it may be possible to supplement statistical evaluations with correlation statistics adapted to non-monotonic relationships, although this type of relationship makes interpretation extremely difficult. In addition, although the present work is based on the explanations provided by *SHAP* (*TreeExplainer*), it applies to all local explanation techniques that make it possible to dissociate the main effect from that of interactions such as complete or coalitionnal methods [6]. More than just a way of rendering information, such a methodology could be applied to many biomedical contexts, including RNA-sequencing, enabling us to improve our understanding of pathophysiology.

## Declarations

## Ethics approval and consent to participate

Not applicable

## Consent for publication

Not applicable

## Data availability

All required data are available at project home page and more details can be found at [8].

## Availability of source code and requirements

– Project name: Explanations interactions
– Project home page: https://github.com/paulmonsarrat/explanations_interactions
– Operating system(s): Platform independent
– Programming language: Python
– License: GNU GPL

## Author contribution

Conceptualization by F.F., J.A., P.M. and L.C. Data curation by F.F. and P.M. Formal analysis by F.F., M.T., E.D., L.C. and P.M. Methodology by F.F., J.A., C.D., E.D. and P.M. Project administration by L.C. and P.M. Software by F.F. Visualization by F.F., C.D., L.C. and P.M. Writing—original draft by F.F. and P.M. Writing—review & editing by J.A., M.T., C.D. and L.C. Resources by L.C. Supervision by J.A., L.C. and P.M. Funding acquisition by J.A., L.C. and P.M.

